# Sensitivity and specificity of human point-of-care circulating cathodic antigen (POC-CCA) test in African livestock for rapid diagnosis of schistosomiasis: a Bayesian latent class analysis

**DOI:** 10.1101/2022.08.17.504243

**Authors:** Beatriz Calvo-Urbano, Elsa Léger, Isobel Gabain, Claudia J. De Dood, Nicolas D. Diouf, Anna Borlase, James W. Rudge, Paul L. A. M. Corstjens, Mariama Sène, Govert J. Van Dam, Martin Walker, Joanne P. Webster

**Author notes:** Corresponding authors: Beatriz Calvo-Urbano^1^, Joanne P. Webster.

## Abstract

Schistosomiasis is a major neglected tropical disease (NTD) affecting both humans and animals. The morbidity and mortality inflicted upon livestock in sub-Saharan Africa has been largely overlooked, in part due to a lack of validated sensitive and specific tests, which do not require specialist training or equipment to deliver and interpret. Inexpensive, non-invasive, and sensitive diagnostic tests for livestock-use would also facilitate both prevalence mapping and appropriate intervention programmes. The aim of this study was to assess the sensitivity and specificity of the currently available point-of-care circulating cathodic antigen test (POC-CCA), designed for *Schistosoma mansoni* detection in humans, for the detection of intestinal livestock schistosomiasis caused by *Schistosoma bovis* and *Schistosoma curassoni*. POC-CCA, together with the circulating anodic antigen (CAA) test, miracidial hatching technique (MHT) and organ and mesentery inspection (for animals from abattoirs only), were applied to samples collected from 195 animals (56 cattle and 139 small ruminants (goats and sheep) from abattoirs and living populations) from Senegal. POC-CCA sensitivity varied by ruminant group and by location/parasite species: sensitivity was greater in Barkedji (cattle: mean 81% (95% credible interval (CrI): 55%-98%); small ruminants: 49% (29%-87%), where livestock were primarily infected by *S. curassoni*, than in Richard Toll (cattle: 62% (41%-84%); small ruminants: 12% (1%-37%), where *S. bovis* was the main parasite species. Mean POC-CCA specificity across sites in small ruminants was 91% (77%-99%) with little variation between locations/parasites (Barkedji: 91% (73%-99%); Richard Toll: 88% (65% - 99%). Specificity could not be assessed in cattle owing to the low number of uninfected cattle surveyed. Overall, our results indicate that, whilst the current POC-CCA does represent a potential diagnostic tool for animal schistosomiasis, future work is needed to develop a livestock-specific affordable and field-applicable diagnostic tests to enable determination of the true extent of livestock schistosomiasis.

**Author summary:** Schistosomiasis is a debilitating neglected tropical and zoonotic disease, infecting over 230 million people and multiple millions of animals worldwide, most notably amongst the poorest regions and populations. The potential contribution of livestock schistosomiasis to disease transmission in human populations has implications for the design of effective disease management and elimination programmes. However, our understanding of the true prevalence, transmission and impact of animal schistosomiasis is severely limited, in part due to a lack of inexpensive, accessible, sensitive and specific diagnostic tools. As a point-of-care circulating cathodic antigen (POC-CCA) diagnostic test is now in widespread use to assess intestinal schistosomiasis caused by *Schistosoma mansoni* in humans, we hypothesised that the same test could be used to detect livestock intestinal schistosomiasis caused by *Schistosoma bovis* and *Schistosoma curassoni*. The aim of this study was thus to evaluate the sensitivity and specificity of the POC-CCA for the detection of intestinal livestock schistosomiasis in Senegal. POC-CCA sensitivity varied by ruminant group and by location/parasite species, while POC-CCA specificity in small ruminants, at least, did not vary across sites. We conclude that the currently-available POC-CCA does represent a potential diagnostic tool for animal schistosomiasis, but that the factors determining test performance warrant further investigation.

## Introduction

The development and application of sensitive and specific diagnostic techniques for detection of infectious diseases is vital for the monitoring and evaluation of all treatment programs in endemic areas, especially whenever considering elimination and/or drug-resistance pharmacovigilance. Within this, point-of-care diagnostic testing is particularly needed wherever there is a necessity for a fast diagnostic outcome that is independent from sophisticated, time-consuming, labour-intense and/or expensive laboratory procedures [1]. This need may be most exemplified for the neglected tropical and zoonotic diseases (NTDs/NZDs), as clearly stressed within the new World Health Organization’s (WHO) NTD Roadmap for 2021-2030 [2] and revised WHO Guideline for the control and elimination of human schistosomiasis, most notably that of recommendation 6 [3, 4].

Schistosomiasis is one of the major debilitating NTDs/NZDs, caused by snail-borne dioecious *Schistosoma* blood-flukes. Approximately 90% of the 230 million people infected worldwide live in sub-Saharan Africa [2]. Animal schistosomiasis is also of major veterinary and socio-economic importance, causing significant mortality and morbidity to livestock, as well as reduced productivity for their owners, although the contribution and consequences of this within sub-Saharan Africa are only just beginning to be realised [2, 5–8]. The main *Schistosoma* species found amongst livestock in Africa are the intestinal *Schistosoma bovis* and *Schistosoma curassoni* in West Africa and *Schistosoma mattheei* in East Africa. These species are phylogenetically close to the human urogenital parasite *Schistosoma haematobium* and have been found to regularly form viable hybrids within humans, both across Africa [8–13], and even within its recent expanse to Europe [14, 15]. Outside of Asia, the contribution of zoonotic schistosomes to human schistosomiasis cases have been largely ignored, despite prevalence levels in humans often remaining unacceptably high following high coverage mass drug administration (MDA) programmes across much of West Africa in particular [8, 16, 17]. Furthermore, recent work combining epidemiological, molecular and mathematical modelling work has demonstrated that the relative role of zoonotic transmission from livestock in Africa is likely to increase as disease control efforts move towards elimination [18].

The newly launched WHO NTD 2021-2030 Roadmap and Guideline for the control and elimination of human schistosomiasis therefore poses the question of anti-schistosomiasis treatment of livestock across Africa in order to achieve the new targets of Elimination as a Public Health Problem (EPHP) in all 78 currently-endemic counties and Interruption of Transmission (IoT) in selected African regions by 2030 [2, 3]. Widespread, indiscriminate use of anthelmintics in livestock have increased drug-resistance [19, 20]. This is particularly pertinent for schistosomiasis as there is currently only one efficacious drug, praziquantel (PZQ), available for both humans and livestock. Recent work has highlighted examples of use and misuse of PZQ in livestock, including, but not exclusive to, PZQ tablets donated free via the WHO MDA programmes for school-aged children being used instead for infected livestock, with little knowledge of application or dosage requirements [5, 7]. Furthermore, recent surveys and socio-economic analyses have revealed highly important, if also often overlooked, animal welfare, productivity, and financial impacts of animal schistosomiasis, which further jeopardize livelihoods, food security and nutrition amongst neglected communities [5]. Indeed the financial costs incurred to subsistence farmers of infected animals was projected to be significantly greater than would be those of a theoretical test and treat programme [5].

To promote sustainable livestock schistosomiasis prevention and treatment in sub-Saharan Africa (SSA), there is therefore a need to both easily and inexpensively diagnose animal schistosomiasis, and subsequently effectively treat only infected individuals and/or herds, such as through a targeted test-and-treat (TnT) or *T3*: *Test*, *Treat*, *Track* design for livestock schistosomiasis in SSA, which anchors the key recent WHO policy recommendations on diagnostic testing, treatment and surveillance in general.

There are, however, currently limited diagnostic techniques that can detect schistosomiasis in livestock with sufficient levels of sensitivity and specificity, as well as logistical ease, that can fully inform disease management decisions [21]. Of those available, a recent systematic review (although data analysed were exclusive to *Schistosoma mansoni* and *Schistosoma japonicum* infections) has recommended the formalin-ethyl acetate sedimentation-digestion with quantitative polymerase chain reaction, as the most promising field-applicable techniques in non-human animal hosts [22], although both are time consuming and/or expensive. A recent extensive field survey of both abattoir and live-sampled livestock within Senegal found some utility with both the Kato-Katz technique and miracidial hatching technique (MHT), but the latter was again labour-intensive and showed significant differences in sensitivities by host and/or parasite species [8].

Tests for antigens, rather than antibodies, are preferred in endemic areas due to the lag in clearance of parasite-specific antibodies after infection subsides [23] and therefore immunochromatographic circulating cathodic antigen (CCA) and circulating anodic antigen (CAA) tests for adult worm antigens in urine (or serum) were developed specifically for current *Schistosoma* infections amongst humans [24]. Whilst a high sensitivity laboratory-based lateral flow (LF) test platform utilizing luminescent up-converting reporter particles (UCP) comprises assay formats for quantitative detection of circulating cathodic and anodic antigen detection in urine (respectively, UCCA and UCAA assays) is available [25, 26], rapid point-of-care testing with visual detection is currently only available for the CCA (POC-CCA, Rapid Medical Diagnostics, Pretoria, South Africa) [27, 28]. Given its utility and field-friendly application, this semi-quantitative (negative, trace, “single positive” +, “double positive” ++ or “triple positive” +++) POC-CCA is now recommended by the WHO for mapping human *S. mansoni* prevalence in endemic areas [29, 30]. The application for detection of human urogenital *S. haematobium* infections with POC-CCA is less efficient i.e. to find the lower worm burden infections [31–34].

The POC-CCA has been used to assess *S. mansoni* infection in one non-human-primate study [35]). Given that *S. bovis* and *S. curassoni* (as well as *S. mattheei*) are intestinal schistosomes of livestock, we predicted that the current POC-CCA could also provide a useful rapid and inexpensive diagnostic for animal intestinal schistosomiasis, as it does for human intestinal schistosomiasis. The aim of this study was therefore to evaluate the sensitivity and specificity of the currently available, human-focused, POC-CCA for the detection of intestinal livestock schistosomiasis caused by *S. bovis, S. curassoni* and their hybrids in Senegal, by host species, in relation to traditional and novel alternative parasitological and immunological diagnostic methods currently available, employing Bayesian latent class models.

## Methods

### Study design and sites

Cross-sectional livestock parasitological surveys were conducted from May to August 2016 and from October 2017 to January 2018 in two areas in Northern Senegal, West Africa; the town of Richard Toll, in the Senegal River Basin, and villages around the Lac de Guiers (area hereafter referred to as Richard Toll), and the town of Linguere and villages around Barkedji along the Vallée du Ferlo (area hereafter referred to as Barkedji). Following the construction of the Diama Dam in 1986, situated on the Senegal River over 100 km downstream from Richard Toll, the environment surrounding this location has undergone important permanent alterations. Desalination, creation of irrigation canals and creation of permanent freshwater bodies have facilitated the expansion of *Schistosoma* snail intermediate host and human-livestock water contact points throughout the year, supporting the co-occurrence and interspecific interactions between *S. haematobium, S. mansoni*, and other *Schistosoma* spp. of veterinary importance [36–38]. The main livestock schistosome species circulating in Richard Toll is *S. bovis* and cattle is the most affected host species. The Barkedji area presents temporary water sources that disappear during the dry season, leading to important seasonal migration of livestock-keeping communities, and annual interruption in schistosomiasis transmission. Small ruminants are the most important livestock host species, infected by *S. curassoni*. However, both schistosome species and hybrids are present in the two areas and all livestock species can be found infected to a lesser extent. Full details on the infection prevalence amongst definitive and intermediate hosts, together with the habitat of two areas, can be found in [8].

Animal sampling was part of a larger survey conducted in these two areas [8]. All animals (cattle, sheep, and goats) routinely slaughtered as part of the normal work of the abattoirs and available for inspection at the time of the surveys were examined post-mortem. Living animals were randomly selected in each village with the initial randomisation carried out at the unit level (owner in this case) and a maximum of five animals of each species (cattle, sheep and goats) then randomly sampled from each selected owner (fewer if the owner had fewer than five animals). Randomisation was carried out using random number generators.

### Diagnostic methods

Whilst adult worms can only be extracted from dead animal (e.g. abattoir or culled animals), infectious status of live animals can be assessed indirectly through parasitological, molecular or immunological methods. Direct and indirect assessment were carried out post-mortem at slaughterhouses and indirect evaluations were applied ante-mortem within the community livestock (see S1). The mesenteric vessels of slaughtered animals were visually inspected for *Schistosoma* adult worms (single males, single females, and paired worms) and those found were stored in RNA-later for molecular analysis (see S2). Faecal, urine, lung, and liver samples were collected post-mortem and were examined for infection using the miracidial hatching technique (MHT) (see S1). Free-swimming miracidia were individually pipetted onto Whatman FTA cards (GE Healthcare Life Sciences, UK) for deoxyribonucleic acid (DNA) storage and subsequent genotyping (see S2). Only faecal and urine samples were obtained from live animals. Faecal samples were assessed for the presence of *Schistosoma* eggs via two Kato-Katz (KK) slides and MHT (see S1 and S2). Animals for which a sufficient volume of urine (≥15 mL) was collected were tested on-site for schistosomiasis with a single point of care-circulating cathodic antigen (POC-CCA) cassette (Rapid Medical Diagnostics, Pretoria, South Africa) and for haematuria with a single Hemastix strip (Siemens Healthcare Diagnostics, Surrey, UK) (see S1). The remaining urine was frozen and transported to Leiden Medical University in the Netherlands for application with the up-converting phosphor-lateral flow (UCP-LF) based assays formats (UCCA and UCAA) for the detection and quantitation of circulating cathodic and anodic antigen in urine (see S1) [39]. Definitions of positive results are described in Table S1.

### Statistical analyses

#### Association between haematuria and POC-CCA results

When extracting urine from abattoir animals, some samples became contaminated with blood. As false-positive POC-CCA tests in humans have been associated with haematuria [40], it was of interest to assess whether POC-CCA results were affected by the blood in urine. Hemastix results were dichotomised as negative (score 0) or positive (scores 1 and 2) and by means of logistic regression we tested the hypothesis that POC-CCA results were independent of blood-contamination status.

#### Composite reference standard and Bayesian latent class model specification

Due to the absence of a gold standard for the diagnosis of schistosomiasis against which we could evaluate the performance of POC-CCA, we derived a composite reference standard (CRS) that was based on three additional diagnostic methods, namely MHT, KK and UCAA. CRS is based on a combination of tests with moderate sensitivity and high specificity [41]. CRS results were considered positive if animals tested positive for either UCAA, MHT or KK. CRS results were assumed to be negative if animals tested negative for all three tests.

We then developed a Bayesian latent class model (BLCM) to assess the sensitivity and specificity of POC-CCA for the detection of active schistosomiasis in cattle and small ruminants [42–44] that took account of the imperfect reference diagnostic tests employed. Two latent (i.e. non-observed) classes were assumed, corresponding to either an infected or non-infected status, and these classes were related to the outcomes of POC-CCA, CRS and UCCA by means of a multinomial distribution [42, 45, 46]. We developed a two-test model and a three-test model. The diagnostic tests included in the two-test model were POC-CCA and CRS. The three-test model made use of the UCCA results and comprised POC-CCA, CRS and UCCA outcomes, albeit at the expense of lower sample sizes in each combination of diagnostic tests.

#### Accuracy model assumptions

Latent class models (LCM), as proposed by Hui and Walter [43], involve three assumptions. Firstly, more than one population needs to be assessed, each with distinct disease prevalence. Secondly, diagnostic test accuracy must remain constant across populations. Thirdly, the accuracy of the tests must be conditionally independent, so that the sensitivity or specificity of one test is independent of the results of a second test [42, 45, 46]. In the present study we were planning to model abattoir and live animals as two distinct populations. However, as schistosomiasis observed prevalence in these two populations were similar (see Table S4), it was likely that the assumption of different prevalences amongst populations was not satisfied. Hence, abattoir and live data were combined and a one population approach was adopted for each site and host species group [44]. However, and in order to assess whether test accuracy differed in these two populations, independent abattoir and live estimates were subsequently derived and compared. To help overcome potential identifiability problems that a one population approach could entail and taking into account the assumed high specificity and medium sensitivity of CRS, we employed moderately informative priors for the CRS (see Table S3). Equally, prevalence priors were moderately informative and their values were based on results from a study carried out in the same area [8]. The impact that these moderately informative priors had on the accuracy estimates (sensitivity/specificity) was assessed through a sensitivity analysis (see S6).

Conditional independence of POC-CCA and CRS in the two-test model was assessed in two ways. Firstly, the Deviance Information Criterion (DIC) [47] of model variants that included (dependence) and excluded (independence) covariance terms were estimated and compared. Models with lower DIC were preferred over models with higher DIC. Secondly, 95% CrI of covariance parameters were assessed [46], with CrIs intersecting zero indicative of conditional independence. Three-test models included dependence terms for POC-CCA and UCCA only as the two tests measure the same antigen (CCA) and their outcomes may not be independent of each other. Furthermore, the covariances between POC-CCA and CRS had been found not to be relevant (see results section “POC-CCA accuracy”).

#### Selection of priors and priors’ sensitivity analyses

The beta distributions used to model the multinomial distribution parameters (Table S3) were parameterized by specifying the mode and the minimum/maximum accuracy value that was believed to be true for each variable, with 95% certainty. Priors for UCAA and UCCA sensitivity and specificity were established based on expert knowledge and published records of high performance [48–51]. Priors for POC-CCA accuracy were non-informative beta (1,1) distributions. Prevalence priors were based on post-mortem examination of abattoir specimens from a previous study [8], where 81% and 82% of cattle were found to be infected in Barkedji and Richard Toll, respectively, and 26% and 16% of small ruminants were found to be infected in Barkedji and Richard Toll, respectively. As the proportion of infected animals were similar at the two sites, only one prior distribution per ruminant group was defined (Table S3). The sensitivity of the results to the prior parameterisations was undertaken by re-estimating POC-CCA accuracy assuming more diffuse priors for prevalence and CRS accuracy (see S6).

#### Accuracy comparisons between sites

In order to assess whether test accuracy differed between sites, differences in the posterior distributions of accuracy between sites were calculated and the probability that the accuracy in one location was greater than in the other was calculated (the Bayesian p-value) by means of the JAGS “step” function [52, 53]. When accuracy did not differ between sites, combined accuracy values were calculated.

#### Software

Statistical analyses were carried out in R (BC7) version 4.0.5 (see S3)

### Ethics approval and consent to participate

For all primary data collection activities, the researchers first explained what the study was about, how the data collection would work and the rights of the participants. Following that, each participant was asked to give their written consent. Ethical approval was sought and granted by i) the Clinical Research and Ethical Review Board at the Royal Veterinary College; approval number URN 2019 1899-3, and (ii) the Comité National d’Ethique pour la Recherche en Santé (Dakar, Senegal) approval number SEN15/68 and SEN 19/68. Urine samples were imported from Senegal to the RVC in the UK with the import licence ITIMP19.0161 and from the UK to LUMC in the Netherlands with the import licence VGM_IN17-1416-GvW.

## Results

### Descriptive statistics

Abattoir surveys were carried out on 89 animals routinely slaughtered, whilst live animal samples were obtained from 106 cattle, sheep and goats from communities within the study area. Given the low number of goats and sheep that were infected in the present study, their data were grouped and analysed as “small ruminants” (see Table 1). POC-CCA diagnostic results were obtained for all 195 animals, 56 of which were cattle and 139 were small ruminants. The distributions of samples across sites and sources of animals (live or abattoir) are shown in Table 1. Barkedji livestock was primarily infected by *S. curassoni*, whilst animals from Richard Toll shed mainly *S. bovis* eggs (see Table S5).

**Table 1.**
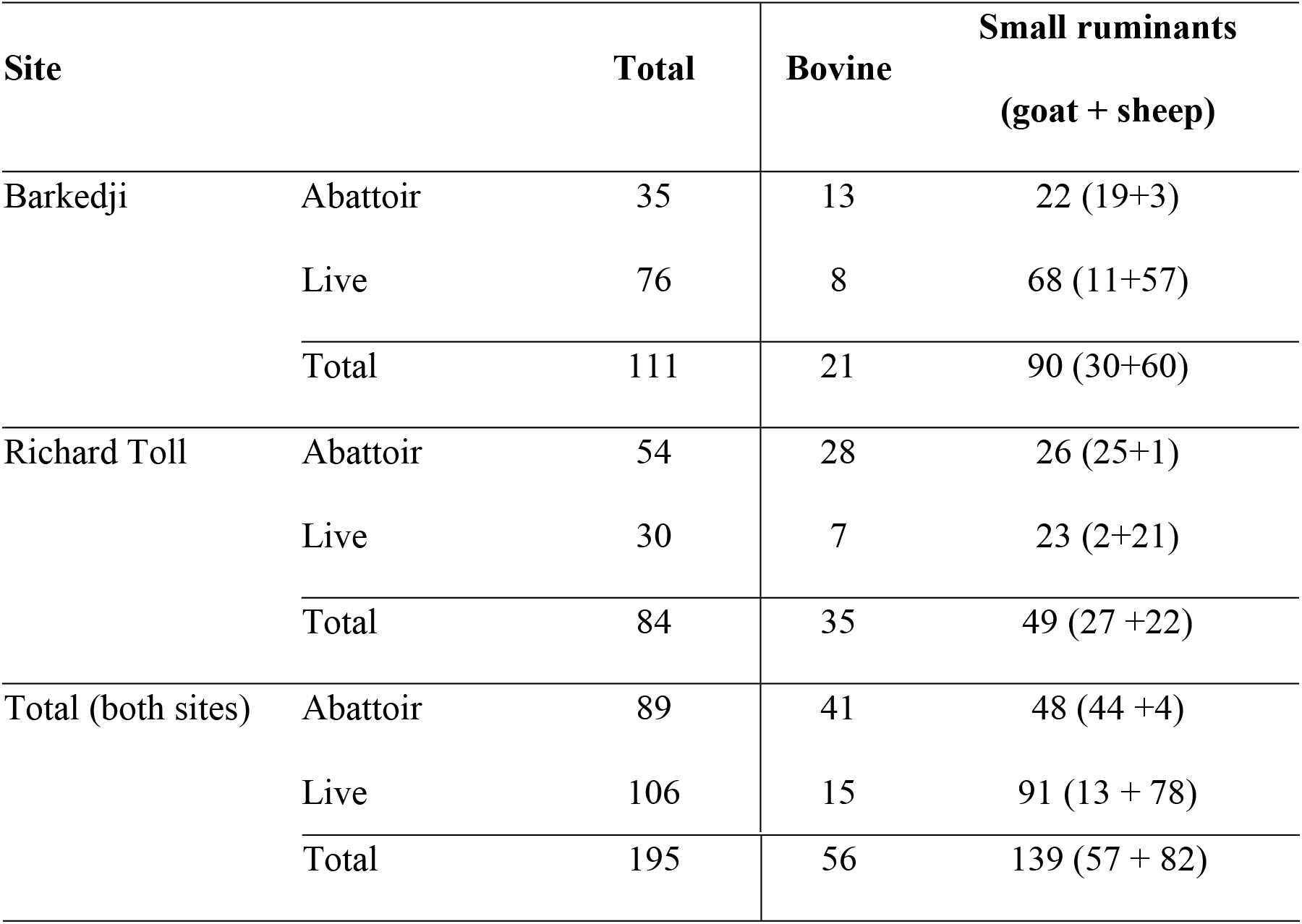
Number of live and abattoir livestock animals surveyed in Barkedji and Richard Toll.

### Association between haematuria and POC-CCA results

Logistic regression results indicated that abattoir POC-CCA results did not depend on Hemastix outcomes, either in cattle (coefficient z value: -0.36, p-value: 0.719) nor in small ruminants (coefficient z value: 0.773, p-value: 0.444). Consequently, in all subsequent analyses, no distinction was made between blood-positive or negative samples.

### Cross-tabulated results

The cross-tabulated results for CRS and POC-CCA are shown in Table 2. The number of small ruminants’ discordant pairs (POC-CCA -, CRS +) was relatively large, suggesting that POC-CCA sensitivity in small ruminants might be low. Cattle were CRS negative in 8 out of 56 animals, showing that most animals were infected, which limited our ability to estimate POC-CCA specificity in cattle. The number of (+, +) concordant pairs was greater in cattle than in small ruminants and the number of (-,-) concordant pairs were greater in small ruminants than in cattle, suggesting that the test behaved differently in each ruminant group.

**Table 2.** Cross-tabulated results for POC-CCA and CRS, by site and ruminant group.

|  |  | Number of concordant and discordant pairs<br>(POC-CCA, CRS)* |  |  |  |
| --- | --- | --- | --- | --- | --- |
| Site | Ruminant | (+, +) | (+, -) | (-, +) | (-, -) |
| Barkedji | Cattle | 13 | 1 | 3 | 1 |
|  | Small ruminants | 19 | 6 | 29 | 35 |
| Richard Toll | Cattle | 17 | 2 | 11 | 4 |
|  | Small ruminants | 3 | 2 | 26 | 18 |

\* POC-CCA = point-of-care cathodic circulation antigen, CRS = composite reference standard

### POC-CCA accuracy

#### Conditional dependence between tests and selection of the best fitting models

BLCM results from the two-test dependence and two-test independence models, by site and ruminant group (Table 3), indicate that POC-CCA and CRS are conditionally independent as: 1) the 95% CrIs of the covariance terms in the two-test model intersected zero; and 2) independence models had lower DIC values than the dependence ones. Results from the three-test model indicate that POC-CCA and UCCA sensitivities were conditionally dependent (lower 95% CrI limit was greater than zero) in all cases with the exception of the small ruminants from Richard Toll (where the CrI contained zero) (Tables 3 and 4). By contrast, POC-CCA and UCCA specificities were conditionally independent, their 95% CrI intersecting zero (Tables 3 and 4). Given the DIC of the two-test independence models were the lowest, we adopted these as the best fitting models and the following conclusions on POC-CCA accuracy were derived from them.

**Table 3.**
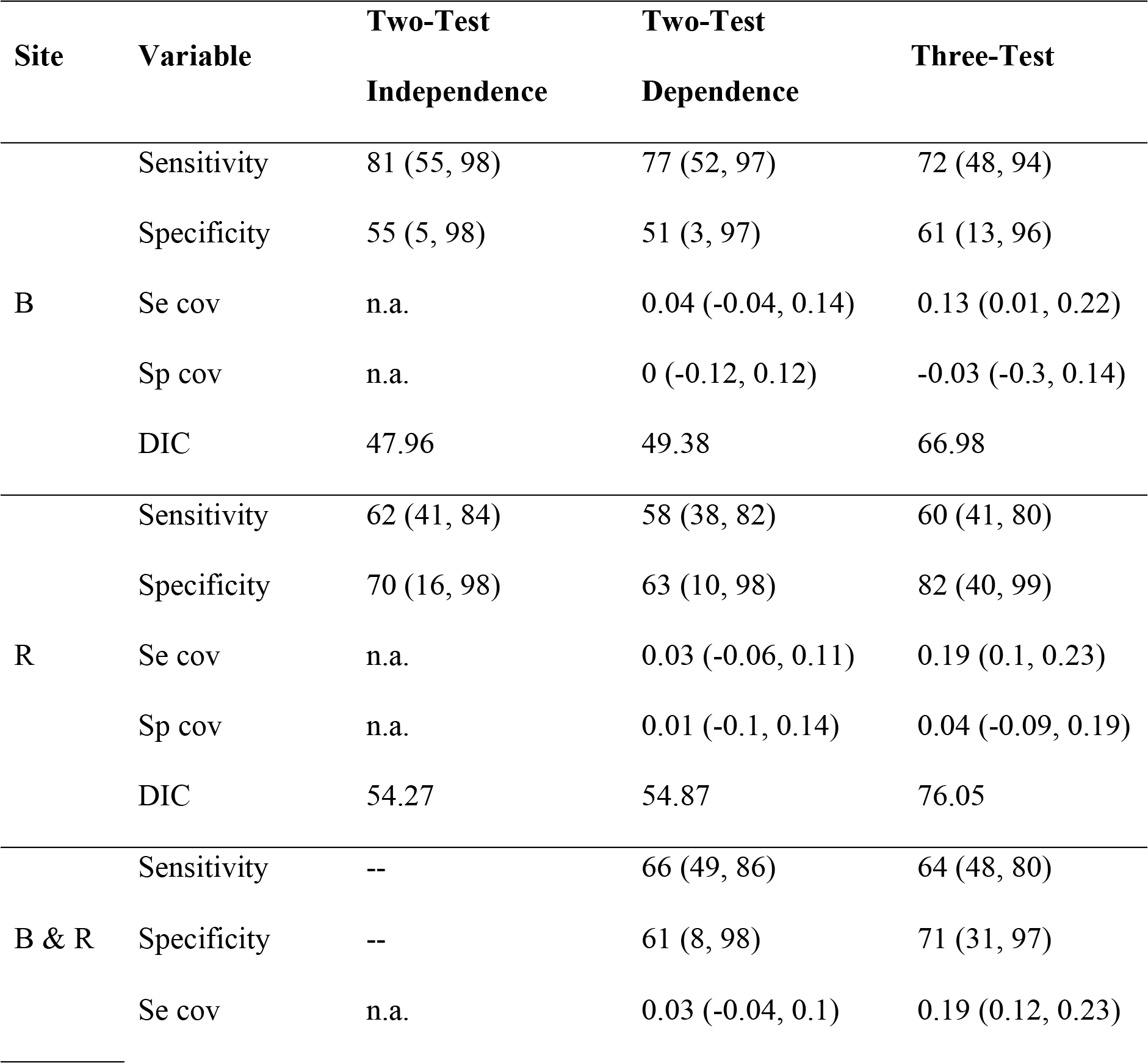
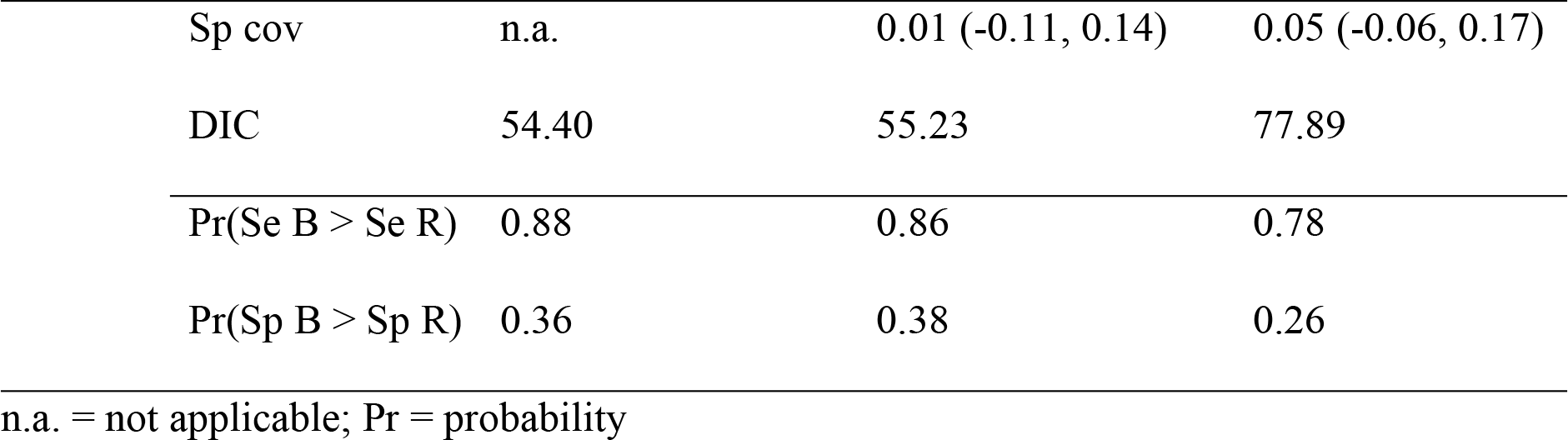
Cattle Bayesian latent class models results from two-test and three-test models showing parameter means (95% CrI) for POC-CCA sensitivity (Se %) and specificity (Sp %) and for covariance (cov), model deviance information criterion (DIC) and Bayesian p-values (Pr) comparing accuracy results between sites. The best fitting model was the two-test independence model. “B & R” rows shows the pooled results for Barkedji (B) and Richard Toll (R).

**Table 4.** Small ruminants Bayesian latent class models results from two-test and three-test models showing parameter means (95% CrI) for POC-CCA sensitivity (Se %) and specificity (Sp %) and for covariance (cov), model deviance information criterion (DIC) and Bayesian p-values (Pr) comparing accuracy results between sites. The best fitting model was the two-test independence model. “B & R” rows shows the pooled results for Barkedji (B) and Richard Toll (R).

| Site | Variable | Two-Test | Two-Test | Three-Tests |
| --- | --- | --- | --- | --- |
|  |  | Independence | Dependence |  |
| B | Sensitivity | 49 (29, 87) | 42 (7, 85) | 41 (25, 66) |
|  | Specificity | 91 (73, 99) | 85 (58, 99) | 91 (75, 99) |
|  | Se cov | n.a. | 0 (-0.1, 0.1) | 0.09 (0, 0.16) |
|  | Sp cov | n.a. | 0.03 (-0.03, 0.17) | 0.02 (-0.03, 0.12) |
|  | DIC | 55.92 | 55.94 | 107.55 |
| R | Sensitivity | 12 (1, 37) | 13 (1, 40) | 26 (8, 68) |
|  | Specificity | 88 (65, 99) | 89 (65, 99) | 93 (74, 100) |
|  | Se cov | n.a. | -0.02 (-0.13, 0.03) | 0.12 (-0.01, 0.21) |
|  | Sp cov | n.a. | 0.02 (-0.03, 0.11) | 0.01 (-0.05, 0.1) |
|  | DIC | 46.53 | 55.92 | 92.92 |
| B<br>&<br>R | Sensitivity | n.a. | 30 (3, 69) | 34 (19, 65) |
|  | Specificity | 91 (77, 99) | 86 (61, 99) | 92 (80, 99) |
|  | Se cov | n.a. | -0.01 (-0.1, 0.07) | 0.12 (0.04, 0.19) |
|  | Sp cov | n.a. | 0.03 (-0.02, 0.16) | 0.02 (-0.01, 0.09) |
|  | DIC | 54.01 | 54.51 | 106.18 |
|  | Pr(Se B > Se R) | 0.99 | 0.91 | 0.80 |
|  | Pr(Sp B > Sp R) | 0.60 | 0.41 | 0.45 |

n.a. = not applicable; Pr = probability

#### POC-CCA accuracy in cattle

POC-CCA sensitivity in Barkedji was 81% (95% CrI: 55% to 98%) and in Richard Toll it was 62% (95% CrI: 41% to 84%). As the probability that the sensitivity in Barkedji was greater than in Richard Toll was high (0.88, see Pr in Table 3), a combined sensitivity value for both sites was not calculated.

This study was not able to estimate POC-CCA specificity in cattle with precision due to the low number of animals that were not infected. This is shown by the wide credible intervals obtained in the analyses (see Table 3).

#### POC-CCA accuracy in small ruminants

POC-CCA sensitivity in Barkedji was 49% (95% CrI: 29% to 87%) and in Richard Toll it was 12% (95% CrI: 1% to 37%) (see Table 4). The probability that the sensitivity in Barkedji was greater than in Richard Toll was 0.99 (see Pr in Table 4). As this difference was large, overall sensitivity across sites was not estimated.

POC-CCA specificity in Barkedji was 91% (95% CrI: 73% to 99%) and in Richard Toll it was 88% (95% CrI: 65% to 99%). The probability that the specificity in Barkedji was greater than in Richard Toll was 0.60. The overall specificity was 91% (95% CrI: 77% to 99%).

#### Prior sensitivity analysis and comparison of accuracy in abattoir and live populations

The results of the prior sensitivity analyses indicate that the baseline prior models adopted were robust and that the priors were not exerting an unduly effect on the POC-CCA accuracy estimates (see S6).

Mean POC-CCA accuracy in abattoir and live populations were similar (not shown), although their 95% CrI were wider than those of pooled samples.

## Discussion

In multi-host, multi-parasite systems, all reservoir hosts must be considered in order to achieve disease elimination. Accurate detection of *Schistosoma* infection in animals would provide not only critical information to guide surveillance and inform control [2–4], but also help to improve livestock-keeping communities wellbeing, finances, and animal welfare [5, 7]. As WHO therefore calls for consideration of the need to treat livestock within Africa in order to minimize zoonotic transmission to humans, as well as for improved diagnostics in general [2–4], accurate diagnosis of schistosomiasis at both the individual and population levels is required for sustainable control programmes as well as assessing, and mitigating against, changes in drug efficacy. This study thereby evaluated the clinical performance of the commercially-available POC-CCA, a diagnostic test routinely used for the detection of the human intestinal parasite, *S. mansoni*, for intestinal *S. bovis, S. curassoni* and hybridized schistosomiasis infections within ruminant livestock of Senegal, West Africa.

We compared the accuracy results obtained in this study with those of the routinely-employed diagnostic techniques, MHT and KK, reported from Senegal by ruminant and parasite species [8], and found that POC-CCA sensitivity was better than that of MHT and KK in Barkedji, where *S. curassoni* was the main parasite species, and similar or inferior in Richard Toll, where *S. bovis* was the main parasite. We are not aware of other studies in the literature reporting accuracy results for the detection of CCA in *S. bovis* or *S. curassoni* infections in livestock. However, CCA has been reported as being strongly associated with *S. bovis* worm and faecal egg counts in goats [54, 55] and CAA were detected in *S. mattheei* infected cows [56].

The test manufacturers have found POC-CCA accuracy to vary by parasite species, indicating that the test was particularly useful for the detection of human intestinal schistosomiasis caused by *S. mansoni*, and less so in the diagnosis of *S. haematobium* [57]. Kittur *et al*. suggested that *S. haematobium* and *S. mansoni* worms may produce different amounts of CCA or that *S. haematobium* may metabolise it more efficiently [58]. It is plausible that *S. bovis* and *S. curassoni* similarly have different patterns of CCA excretion and metabolic pathways, and thus respond differently in POC-CCA accuracy. For instance, if the maximum number of adult worms a host can carry falls beneath the test’s minimum level of detection, infected hosts would be misdiagnosed as uninfected. These maxima may differ with parasite species, resulting in differential misclassification. To our knowledge, the respective maximum number of worms each host can carry, by parasite species and ruminant group, has not been determined. The test manufacturers have also found that, in the case of *S. haematobium*, CCA levels vary by region. Moreover, a study comparing POC-CCA performance across five countries found that *S. mansoni* results varied between countries [29]. The present study could not discern between the effects of location and parasite species, as *S. curassoni* was the predominant parasite in Barkedji whilst *S. bovis* was primarily present in Richard Toll. Hence, differences in test accuracy between the two locations could have arisen due to differences in parasite species/hybrid distributions, as well as local factors determining intensity of infection such as intermediate snail distributions. The disparity between each host species’ test performance might be attributable to the factors such as consistency of urine, rate of metabolising CCA, or cross-reaction with other infections.

Our study also found that there were no differences in POC-CCA specificity in small ruminants across sites and parasite species, and that POC-CCA specificity was not affected by haematuria. This contrasts with results in humans, where POC-CCA specificity has been found to be affected by the host’s age (in pre-school aged children < 5 years), their pregnancy status, and whether or not they have haematuria or a urogenital infection [40, 59].

POC-CCA accuracy has been found to vary depending on production batch, raising questions regarding production quality control and calls for the optimisation and standardisation of production [60–62]. This is of particular relevance if POC-CCA is to become reliable a tool with which to determine whether to treat human and livestock populations [2–4].

### Limitations

Due to the absence of a gold-stand diagnostic test for schistosomiasis, this study derived a composite reference standard (CRS) that was based on KK, MHT and UCAA results. The sensitivities of KK and MHT are relatively low, partly due to the technical difficulties associated with the management of large volumes of faecal material. These methods are highly specific, although misclassification of eggs and miracidia can occur. On the other hand, UCAA assays had been labelled as ultrasensitive for human schistosomiasis detection [51, 63], but have not been optimised for livestock. By combining these tests into a CRS we aimed to obtain a composite measure that was highly specific and moderately sensitive. However, we found eight cases where UCAA was negative and KK or MHT were positive, indicating that the high sensitivity UCAA test may not detect all the infections despite its lower limit of detection of 0.6 pg/mL CAA [63]. Given the variable quality of the urine samples, it was unfortunate that paired serum samples were not available to validate urine testing. Following recommendations in the literature for human schistosomiasis and in order to identify the most infections [64], the present study considered POC-CCA trace as a positive result. The relatively low sample sizes for goats, sheep and infected cattle had an impact on the precision (wide 95% CrI) of our estimates and our ability to analyse goat and sheep data separately. Lastly, we were not able to assess the interactions between location, ruminant group and parasite species. It is likely that POC-CCA accuracy in Barkedji and Richard Toll can be assimilated to accuracy for *S. curassoni* and *S. bovis* detection, respectively. However, this need to be further investigated. The observed high probabilities of sensitivity differences point to novel evidence of variation between host and parasite spp. that can inform the design and analyses of future diagnostic evaluations.

### Implications for practice

Overall, our results indicate that POC-CCA represents a potential diagnostic tool for schistosomiasis in ruminant livestock populations. However, in order to move towards the interruption of transmission, the elimination of this zoonotic transmitted disease [2, 3] and to safeguard the welfare of livestock and the livelihoods of the communities that depend on them [5], it is of great importance to develop inexpensive livestock-specific POC-CCA tests that enable us to formulate accurate assessments of disease prevalence. Furthermore, the observed variation in test performance across sites and parasite species has implications for the applicability of this diagnostic method, as it may hinder our ability to establish universally valid thresholds for disease prevalence that inform control programmes. Hence, the factors that determine test performance need to be investigated further so that region-specific guidelines could be derived if needed. Lastly, manufacturers quality control must be a foremost priority if POC-CCA diagnostic tests are implemented for the assessment of schistosomiasis, both in humans and animals.

## Acknowledgments

We are extremely grateful to all the Senegalese communities involved in the project. We are particularly grateful to the late Mr. Cheikh Tidiane Thiam, Mr. Samba Deguene Diop, Mr. Alassane Ndiaye and Mr Stefano Catalano for assistance in the field, and Dr Cheikh B Fall for facilitating transfer of the samples. We are also extremely grateful to Prof David Rollinson of the Natural History Museum and Global Schistosomiasis Alliance for invaluable discussions at the project onset. We also sincerely thank Dr Aidan Emery at the Schistosomiasis Collection at the Natural History Museum (SCAN) for his help on the import of the urine samples from Senegal to the UK.

## Availability of data and materials

All data generated or analysed during this study, together with the coding developed, are included in this published article [and its supplementary information files]. Other datasets used and/or analysed can be made available by the corresponding author on reasonable request.

## Consent for publication

Not applicable.

## Authors’ contributions

JPW conceptualized the study; EL, ND, AB, MS and JPW performed field work and/or facilitated access to farmers; CJDD, PLAMC and GVD performed CAA validation, BC-U performed statistical analyses with input or guidance from IG, AB, MW and JPW. Original draft preparation was performed by BC-U, EL, IG and JPW, with all authors contributing and approving the final manuscript.

## Supporting information Captions

**S1. Diagnostic tests**

**S1 Text. Description of diagnostic techniques**

**S2. Molecular analyses for *Schistosoma* species determination**

**S2 Text. Description of molecular analyses for the determination of *Schistosoma* species**

**S3. Statistical analysis**

**S3 Text. Latent Class Models**

**S3 Table. Baseline priors used in multivariate modelling.**

**S4. Observed prevalence**

**S4 Table. Summary statistics by diagnostic method: number of infected animals/number of animals examined (empirical prevalence), per ruminant group R: C = cattle, SR = small ruminants), site (column S: B = Barkedji, R = Richard Toll) and animal population (column P: A = abattoir, L = live animals).**

**S5. Distribution of *Schistosoma* species**

**S5 Text. Distribution of *Schistosoma* species**

**S5 Table. Number of animals genotyped and number of animals in each combination of *Schistosoma* species.**

**S6. BLCM sensitivity**

**S6.1 Table. Priors adopted in the sensitivity analysis, modifying CRS priors (A) and prevalence priors (B).**

**S6 Text**

**S6.2 Table. BLCM results from 2-tests independence analyses showing parameter means (95% CrI) and models’ deviance information criterium (DIC), for POC-CCA sensitivity (Se, %) and specificity (Sp, %), comparing baseline priors, CRS modified priors and prevalence modified priors.**

**S7. Models JAGS code**

**S7.1 Text. JAGS code for two-test independence/dependence models [20].**

**S7.2 Text. JAGS code for three-test model [21].**

**S 8. Appendix references**

